# Linking gene expression to unilateral pollen-pistil reproductive barriers

**DOI:** 10.1101/080663

**Authors:** Amanda K. Broz, Rafael F. Guerrero, April M. Randle, You Soon Baek, Matthew W. Hahn, Patricia A. Bedinger

**Author notes:** Email addresses of all authors: AKB:; RFG:; AMR:; YSB:; MWH:; PAB. Author for correspondence: Patricia Bedinger. These authors contributed equally to this work (Amanda K. Broz, Rafael F. Guerrero).

## Abstract

Unilateral incompatibility (UI) is an asymmetric reproductive barrier that unidirectionally prevents gene flow between species and/or populations. UI is characterized by a compatible interaction between partners in one direction, but in the reciprocal cross fertilization fails, generally due to pollen tube rejection by the pistil. Although UI has long been observed in crosses between different species, the underlying molecular mechanisms are only beginning to be characterized. The wild tomato relative *Solanum habrochaites* provides a unique study system to investigate the molecular basis of this reproductive barrier, as populations within the species exhibit both interspecific and interpopulation UI. Here we used a transcriptomic approach to identify genes in both pollen and pistil tissues that may be probable key players in UI. We confirmed UI at the pollen-pistil level between a self-incompatible population and a self-compatible population of *S. habrochaites*. A comparison of gene expression between pollinated styles exhibiting the incompatibility response and unpollinated controls revealed only a small number of differentially expressed transcripts. Many more differences in transcript profiles were identified between UI-competent versus UI-compromised reproductive tissues. A number of intriguing candidate genes were highly differentially expressed, including a putative pollen arabinogalactan protein, a stylar Kunitz family protease inhibitor, and a stylar peptide hormone Rapid Alkalinization Factor. Our data also provide transcriptomic evidence that fundamental processes including reactive oxygen species signaling are likely key in UI pollen-pistil interactions between both populations and species. Our transcriptomic analysis highlighted specific genes, including those in ROS signaling pathways that warrant further study in investigations of UI. To our knowledge, this is the first report to identify candidate genes involved in unilateral barriers between populations of the same species.

## Background

Reproductive barriers are critical for maintaining species integrity, and their emergence between populations is associated with decreased gene flow and increased genetic divergence, ultimately leading to speciation. In a number of plant families, including the Solanaceae, a unidirectional post-mating prezygotic barrier termed unilateral incompatibility (UI) occurs at the level of pollen-pistil interactions. UI has most often been studied between genera or species [1–5], but there is also evidence of UI between populations of the same species [1, 6–9].

The unidirectionality of UI in crosses between species has been linked to plant mating system and specifically to the self-incompatibility response. Self-incompatible (SI) species are obligate outcrossers, whereas self-compatible (SC) species are capable of producing offspring through self-pollination. Gametophytic SI is the most common system of incompatibility in flowering plants [10] and in many families it is dependent on the activity of pistil-expressed *S*-locus ribonucleases, or S-RNases [11, 12]. Distinct from the sporophytic SI of the Brassicaseae in which pollen is rejected at the stigma surface [13, 14], in gametophytic SI growing pollen tubes are actively rejected within the style [11, 15–17]. This type of S-RNase-based SI is genetically determined by the polymorphic *S*-locus, which harbors pistil-(S-RNase) and pollen-(S-locus F-box) expressed factors that are required for the specificity of the SI response [18–21]. Additional factors have also been implicated in SI, although they do not determine specificity. These include pistil-expressed HT-proteins and 120K glycoproteins [22–24], as well as pollen-expressed components of the E3 ubiquitin ligase complex (Cullin1 and SKP1) [15, 25–29].

UI generally follows the SI x SC rule, wherein pollen of SC species is unable to fertilize SI ovules, but the reciprocal cross is compatible [4]. Because of this link between SI and UI, initial studies suggested pleiotropic effects of the *S*-locus on interspecific incompatibilities. In fact, more recent studies have shown that the proteins S-RNase, HT-protein, S-locus F-box 23 and Cullin1 all function in both the SI and UI response, demonstrating mechanistic overlap between these two types of reproductive barriers [26, 28, 30–33]. However, it is also clear that genetic factors other than those involved in SI function in UI.

Crosses between SC species or populations (many of which do not express S-RNase) exhibit unexpected incompatibilities [1–4, 7, 31, 34, 35]. This suggests that there are at least two mechanisms underlying UI, one of which is S-RNase based and the other of which is S-RNase independent. Most SC species exhibit full crossability with other SC species; however only recently evolved SC populations of SI species exhibit UI [2, 4]. This suggests that the mechanistic factors underlying S-RNase independent UI are also present in populations that contain S-RNase. However, the number and type of factors involved in UI remains unclear and most models suggest roles for multiple UI genes of both large and small effect [2, 5, 8, 36].

The tomato clade (*Solanum* section *Lycopersicon*) offers ample opportunities to further understand and identify the genes involved in UI [37]. This small clade has recently diverged from a common SI ancestor [38, 39] over the course of ~2.5 million years [40]. The clade harbors six SC species, and seven SI species, three of which contain both SI and SC populations [41]. A number of studies have examined both the physiological aspects [1, 5, 42, 43] and genetic basis [26–28, 30, 31, 33] of UI between members of the clade. In addition, a small number of studies have assessed UI between SI and SC populations within a species [1, 6–9, 43].

The wild tomato *Solanum habrochaites* is an ideal species in which to study both interspecific and interpopulation UI within the context of recently diverged populations. *S. habrochaites* has undergone at least two independent transitions from the ancestral SI state to SC at both the northern and southern species range margins [6,7, 9, 44]. The loss of SI at the northern range margin is correlated with the loss of pistil-side SI factor S-RNase [6, 31, 45]; whereas southern *S. habrochaites* SC populations express an S-RNase protein with little or no RNase activity [31]. The genetic structure of *S. habrochaites* is consistent with the spread of populations both north and south from a central origin, with central SI populations showing the highest levels of diversity [44, 46]. Interestingly, the types and strengths of reproductive barriers displayed by a number of populations also follow this same latitudinal axis of divergence [6, 44].

Previous studies have demonstrated variation in the strength of both intra- and interspecific UI in different *S. habrochaites* populations [1, 6–9, 31, 35]. For instance, a central *S. habrochaites* SI population (LA1777) rejects pollen tubes from a subgroup of northern *S. habrochaites* SC populations, including LA0407; whereas LA0407 and other SC populations accept LA1777 pollen tubes [1, 6].

In interspecific interactions, when either SC *S. lycopersicum* or SC *S. neorickii* (as female) is crossed with *S. habrochaites* (as male), crosses result in fruit [47, 48]. In the reciprocal cross, pistils of SI and southern SC populations of *S. habrochaites* rapidly reject interspecific SC pollen after a few millimeters of growth through the style, demonstrating a classic UI response [1, 6, 31]. However, interspecific pollen tube rejection is delayed in many northern SC populations of *S. habrochaites*, [1, 6, 31], and one unique SC population (LA1223) is unable to reject both *S. lycopersicum* and *S. neorickii* pollen tubes [6]. LA1223 is the only known *S. habrochaites* population that does not accumulate the pistil HT-protein [6, 31], which may explain this extraordinary lack of UI at the pollen-pistil level.

Although elegant mapping, biochemical and transgenic studies have revealed that SI factors are important players in UI [12, 23, 26, 28, 30, 31, 33, 49], the full suite of molecular factors involved in the complex UI response are only beginning to be characterized. The evidence for both interspecific and interpopulation UI in *S. habrochaites* suggests that factors other than those involved in SI are involved in UI. This is consistent with transgenic studies in tomato showing that together S-RNase and HT-protein can act to reject pollen from only a subset SC species, and even then pollen tube rejection is delayed [33].

Transcriptome profiling can provide a broad view of the factors involved in pollen-pistil interactions and represents a powerful tool for candidate gene identification. Studies in *Arabidopsis* have identified large numbers of genes that are specific to pollen or pistil, as well as those that are specifically induced upon the interaction between the two in compatible pollinations [50–53]. However, very few studies have investigated pollen-pistil transcriptomes in response to UI. In a recent study, Pease et al. [54] identified a number of interspecific UI candidate genes by comparing stylar transcriptomes of UI-competent *S. pennellii* and UI-deficient *S. lycopersicum*, two species which likely diverged over 2 mya [40]. Their analysis identified a large number of loci that varied in expression between styles of these two species, and highlighted the importance of HT-protein in UI [54].

Here, we used a similar transcriptomics approach to broadly characterize the molecular factors involved in UI in *S. habrochaites* at the level of pollen-pistil interactions. The populations selected for study have recently diverged (< 0.8 mya, [40]), and exhibit differences in both interspecific and interpopulation reproductive barriers. To our knowledge, this is the first report using transcriptomics to identify novel candidate genes associated with reproductive barriers between diverging populations within a species.

## Results

### Pollen-pistil interactions in SI LA1777 and SC LA0407

Using pollen tube growth assays, we confirmed that all LA1777 (SI) individuals used in RNA-seq experiments rejected both self-pollen tubes and those of LA0407, but accepted pollen from other LA1777 individuals (intrapopulation). Further, we confirmed that each LA0407 (SC) individual accepted pollen tubes from LA1777, self, and intrapopulation crosses.

In compatible crosses, pollen tubes generally reach the ovary within 24 hours [1, 31]. However, incompatible crosses may differ in the timing of pollen tube rejection [1]. To identify the post-pollination time-point at which pollen tube rejection occurs in the incompatible cross between LA1777 (female) and LA0407 (male), pistils were pollinated and harvested over a time course. Pistils of LA1777 were actively rejecting pollen of LA0407 at 12 hours post-pollination, and full rejection occurred by 48 hours (Fig. 1). In an attempt to capture changes in gene expression that would correspond to active pollen tube rejection, we chose to harvest stylar tissue 16 hours post-pollination for subsequent RNA-seq experiments. At this point, incompatible pollen tubes will have traversed ~20% of the style length (Fig. 1), whereas compatible pollen tubes will have traversed ~60% of style length [31].

**Figure 1.**
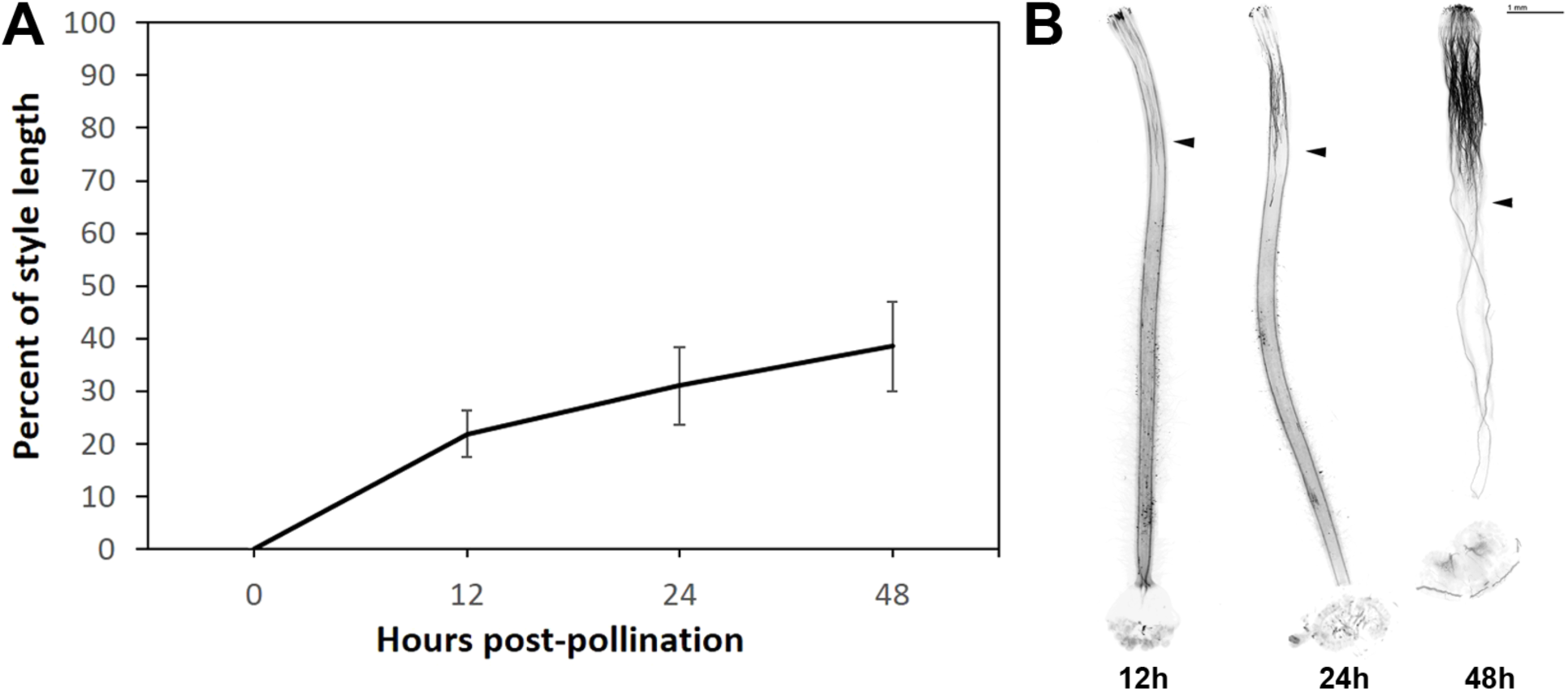
Interpopulation pollen tube growth over time. Pistils of *Solanum habrochaites* LA1777 were pollinated with LA0407 pollen and harvested at 12, 24 and 48 h after pollination. A, Time course of LA0407 pollen tube growth in styles of LA1777 with standard deviation for each time point. B, Representative image of LA0407 pollen tube growth in LA1777 pistils at each time point; arrowhead points to the location where the majority of pollen tubes are rejected.

### Differential gene expression between UI-competent and compromised populations

We sampled both pollen and stylar tissue to better characterize differences in transcriptomes between UI-competent LA1777 and UI-compromised LA0407 (Fig. 2). Styles were either unpollinated (UP) or subject to three different treatments including self, intrapopulation and interpopulation pollination (Fig. 2). We analyzed a total of 33,055 genes, out of which 7,358 (22.3%) were specific to pollen (that is, they were upregulated in pollen with respect to all styles), and 4,793 (14.5%) were specific to styles. Additionally, we found 1,492 genes that were highly upregulated only in tissues of LA1777 and 1,382 genes highly upregulated only in LA0407 (4.5% and 4.2% of genes analyzed, respectively).

**Figure 2.**
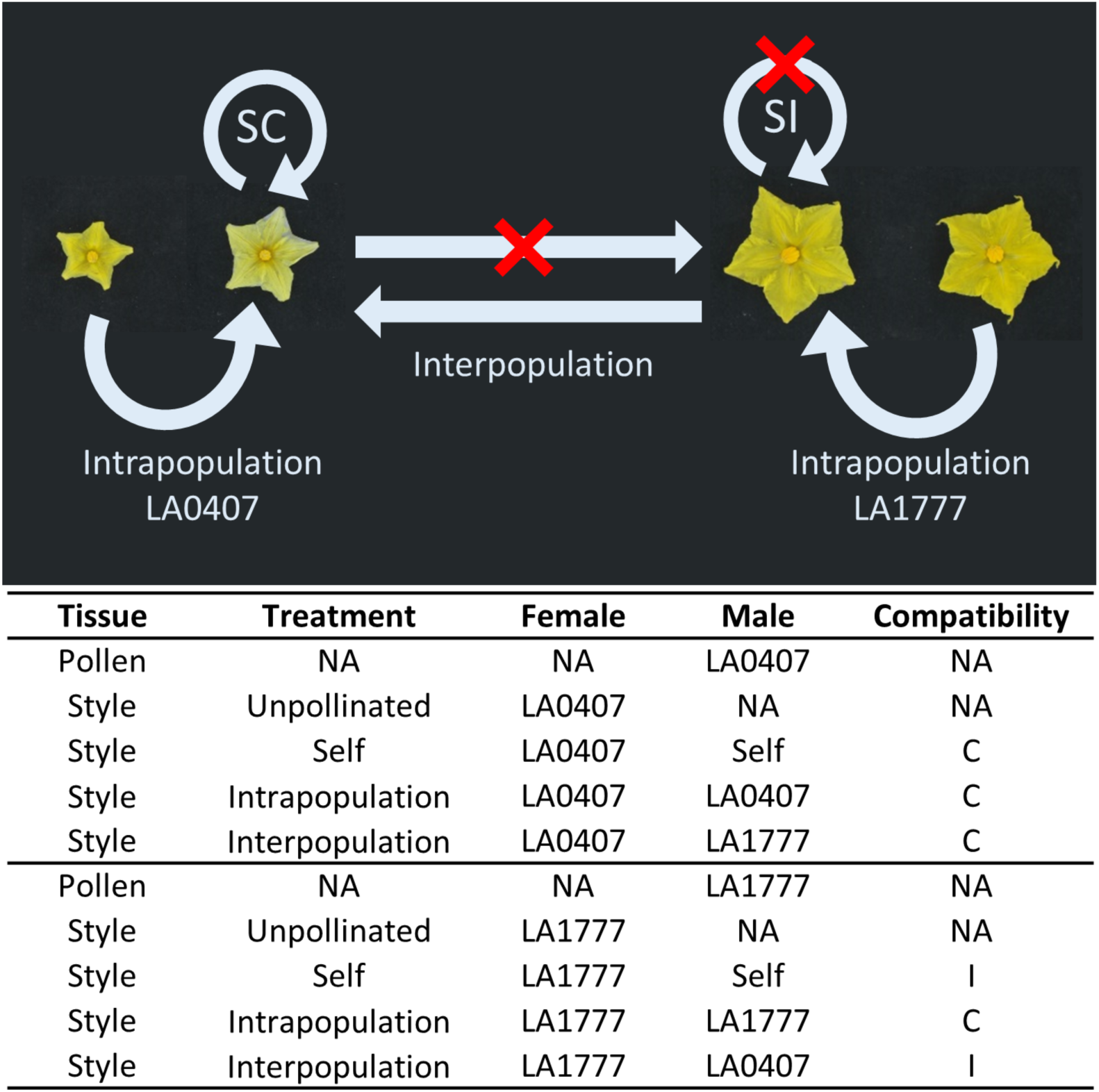
Experimental design for *Solanum habrochaites* RNA-seq experiment. Top panel: compatibilities within and between *S. habrochaites* populations LA0407 and LA1777. SC, self-compatible; SI self-incompatible. Bottom panel: Tissues (pollen, style) and stylar treatment types (intrapopulation, interpopulation, self-pollinations and unpollinated) were collected for three biological replicates of LA0407 and LA1777. C, compatible pollination; I, incompatible pollination; NA, not applicable.

### Pollination type significantly influences a small number of genes

We expected that stylar transcriptomes would exhibit differences due to the type of pollination and whether or not a pollination was compatible. We performed three separate pairwise comparisons to look at differential gene expression in unpollinated (UP) LA1777 styles compared to different pollination types (intrapopulation, self and interpopulation). Pairwise comparisons between pollinated and UP LA1777 styles revealed only a small number of differentially expressed genes, all of which were expressed at very low levels (<0.3 counts per million (cpm)), Additional File 1, Table S1 (self vs UP); Additional File 1 Table S2 (intrapopulation vs UP) and Additional File 1, Table S3 (interpopulation vs UP)). In all pollination treatments we identified pollination-induced differential expression of a number of hypothetical proteins, RNA-directed DNA polymerases and retrovirus-related reverse transcriptases. However, only three genes showed similar regulation among all pollination treatments: a hypothetical protein (Sopen05g010690) and two genes that have high homology to a Copia-like retrotransposon (Sopen01g019140 and Sopen02g008750).

Five genes were similarly differentially expressed in incompatible crosses with LA1777 as female (self and interpopulation crosses, Additional File 1, Table S1 and Table S3) versus compatible crosses (intrapopulation, Additional File 1, Table S2), three of which encoded hypothetical proteins. In addition, a cystathionine beta synthase (CBS) domain-containing protein (Sopen06g019460) was downregulated 3-fold in incompatible crosses. It has been proposed that CBS proteins are redox regulators involved in the modification of cell wall composition, and in *Arabidopsis*, changes in CBS expression can reduce self-fertility [55]. A homolog of *Defective meristem silencing 3* (DMS3, Sopen03g019410), a gene that is involved in silencing and epigenetic modification, was also downregulated 3-fold in incompatible crosses [56].

Thirty two genes that were differentially regulated between interpopulation pollination and UP styles (Additional File 1, Table S3), but not in other treatments (self vs UP, Additional File 1, Table S1 or intrapopulation vs UP, Additional File 1, Table S2). Interesting candidates included a Ras-related GTPase (Sopen04g023040, Rab3), a pollen specific calcium binding annexin (Sopen04g003390), and a K^+^ transporter (Sopen11g006300) were upregulated ~3-fold in interpopulation pollinations. Another candidate gene, upregulated 16-fold in interpopulation pollinations encodes a H_2_O_2_ transporter (Sopen10g033580, PIP2). Each of these genes are potentially involved in reactive oxygen species (ROS) signaling during pollen-pistil interactions [57–61], and their upregulation in interpopulation pollinations may alter ROS signaling, resulting in the dysfunctional pollen tube growth characterized by UI. Further, in interpopulation pollinations, an endonuclease was upregulated over 18-fold, and a DNase over 2.9 fold. Intriguingly, in *Pyrus pyrifolia* and *Papaver rhoeas*, the disruption of ROS signaling in incompatible (self) pollen tubes leads to depolymerization of the actin cytoskeleton and an increase in nuclear DNA degradation [62, 63].

### Differential gene expression related to UI-competence in styles

Although a number of potentially interesting candidates involved in UI were identified from our pairwise comparisons, they were expressed at relatively low levels, often showed only low fold-changes, and had relatively high p-values (p < 0.05). A principal components analysis (PCA) of all stylar treatments showed that most of the variation between samples could be explained by source population, and that there was no consistent grouping of styles due to pollination treatment (Additional File 2, Fig. S1). The fact that few significant differences were detected between UP styles and pollination treatments may reflect results of previous studies indicating that the genes involved in UI-competence are expressed in styles regardless of pollination status [31, 37, 54]. In other words, the UI-competent styles appear ‘primed’ to reject incompatible pollen, as large changes in gene expression are not seen between UP styles and those pollinated with compatible versus incompatible pollen parents. Because of these results, we also performed an analysis in which all stylar treatments within a population (UP, pollinated with self, intrapopulation and interpopulation pollen) were pooled to capture differential expression between UI-competent (LA1777) versus UI-compromised (LA0407) styles.

We identified 179 genes that were significantly upregulated in UI-competent versus compromised styles, and 179 that were significantly downregulated (Additional File 1, Table S4 and Table S5). However, we focused our interest predominantly on genes that showed a mean expression level over 3 normalized cpm and were upregulated over 10-fold in UI-competent versus compromised styles. The top 25 genes showing the largest upregulation in UI-competent styles are shown in Table 1.

**Table 1.** Top-25 upregulated genes in styles of LA1777 vs. LA0407.

| Gene | p value | Mean expression | Fold change | Annotation |
| --- | --- | --- | --- | --- |
| Sopen12g017530.1 | 1.9E-20 | 3.8 | 1451.4 | [HP] Hypothetical protein; Contains DUF pfam0403, heavy metal-associated |
| Sopen02g037430.1 | 4.1E-17 | 4.2 | 163.8 | Cytochrome P450; family 81, subfamily D |
| Sopen05g032070.1 | 3.8E-14 | 9.2 | 154.9 | Chalcone Synthase 2; flavonoid biosynthesis |
| Sopen06g021780.1 | 5.1E-13 | 3.9 | 116.2 | Cytochrome P450; family 71, subfamily B |
| Sopen01g011020.1 | 4.0E-21 | 75.9 | 108.4 | Hypothetical protein |
| Sopen12g006330.1 | 8.4E-15 | 22.2 | 58.6 | 2-Alkenal Reductase; involved in response to oxidative stress |
| Sopen08g021280.1 | 3.4E-11 | 8.4 | 51.8 | Chlorophyll a-b Binding Protein 3C |
| Sopen12g026980.1 | 5.7E-14 | 4.0 | 36.6 | Glutathione S-Transferase; involved in stress response |
| Sopen05g003640.1 | 2.8E-19 | 4.3 | 35.6 | NAD(P)H Dehydrogenase Subunit; in chloroplast, PsbQ-like 2 |
| Sopen09g003470.1 | 6.1E-06 | 10.3 | 32.1 | Threonine Dehydratase; isoleucine biosynthesis, plastid |
| Sopen04g002110.1 | 4.9E-12 | 5.2 | 29.6 | Programmed Cell Death 6-Interacting Protein; endosomal |
| Sopen05g001950.1 | 2.6E-17 | 4.3 | 28.7 | Major facilitator superfamily; oligopeptide transport protein |
| Sopen07g028940.1 | 4.8E-11 | 4.5 | 26.7 | Cytochrome P450; family 72, subfamily A |
| Sopen02g033850.1 | 1.7E-10 | 10.6 | 22.0 | [HP] Hypothetical protein; 80% identity to *Solanum chacoense* RALF-like5 |
| Sopen02g027710.1 | 6.0E-09 | 182.2 | 20.2 | Acidic Endochitinase; involved in defense response |
| Sopen02g022730.1 | 2.5E-11 | 5.4 | 19.6 | Dormancy-Associated Auxin-Repressed small protein |
| Sopen02g021850.1 | 2.2E-11 | 3.4 | 18.9 | Transcription factor; bHLH30-like DNA binding superfamily |
| Sopen12g021490.1 | 7.0E-18 | 10.2 | 17.4 | NAD(P)-linked Oxidoreductase superfamily protein; involved in stress response |
| Sopen10g025030.1 | 3.1E-09 | 4.9 | 14.9 | Dirigent-Protein Like 22; involved in defense response |
| Sopen11g004500.1 | 2.8E-08 | 3.8 | 14.4 | Beta Glucosidase 42 (BGLU42) |
| Sopen12g006340.1 | 6.6E-11 | 5.2 | 14.4 | Transferase, HXXXD-type acyl-transferase family protein |
| Sopen01g047950.1 | 1.6E-04 | 17.7 | 14.0 | ABC Transporter G family member 11-like; cutin or wax export, stress response |
| Sopen10g001450.1 | 8.8E-18 | 3.2 | 13.4 | Hypothetical protein, chloroplast |
| Sopen03g029620.1 | 1.0E-04 | 6.8 | 12.8 | Kunitz family trypsin and protease inhibitor (similar to Miraculin) |
| Sopen05g030780.1 | 3.7E-12 | 18.8 | 12.8 | Chalcone-Flavanone Isomerase family protein |

Gene annotation from Spenn\_v2.0 was enhanced, when possible, for genes noted as hypothetical proteins (marked as [HP]).

Within the top 25 genes showing the highest upregulation in UI-competent versus compromised styles, we found ten that are involved in oxidation-reduction reactions (Table 1). These included three Cytochrome P450 genes (Sopen02g037430, Sopen06g021780, Sopen07g028940), an Alkenal Reductase (Sopen12g006330), a Glutathione S-Transferase (Sopen12g026980), an NAD(P)H-Dehydrogenase (Sopen05g003640), and an NAD(P)-Oxidoreductase (Sopen12g021490). In addition, we identified two genes involved in early steps of flavonoid biosynthesis: Chalcone Synthase 2 (Sopen05g032070) and Chalcone-Flavanone Isomerase (Sopen05g030780). Finally, we discovered that the gene showing the highest upregulation in UI-competent styles (hypothetical protein, Sopen12g017530) contains a heavy metal-associated domain (Additional File 2, Fig. S2), suggesting that it also may be involved in oxidation-reduction reactions.

In UI-competent styles, we also identified three upregulated genes that are putatively involved in defense response. These included an Endo-Chitinase involved in jasmonic acid and ethylene signaling (Sopen02g027710) that was highly expressed (182 cpm) and upregulated over 20-fold, a Beta-Glucosidase (Sopen11g004500) and a Disease Resistance Protein (Sopen10g025030) (Table 1). These genes are of interest, as many molecular components involved in plant-pathogen interactions show significant overlap with those involved in pollen tube growth and guidance [64].

One gene of particular interest that was highly upregulated in UI-competent styles is the hypothetical protein Sopen02g033850 (Table 1), which shows 80% amino acid identity to the peptide hormone Rapid ALkalinization Factor (RALF) from the wild potato, *S. chacoense*. Small secreted peptides including RALFs are involved in a wide variety of plant functions including development and immunity [65], and a pollen-specific RALF from *S. lycopersicum* (SlPRALF) was found to negatively regulate pollen tube elongation *in vitro* [66]. We also identified a protein involved in oligo-peptide transport (Sopen05g001950) which was upregulated >28-fold in UI-competent styles and may be involved in the transport/secretion of peptide hormones such as RALFs.

Another gene of interest that was highly upregulated in UI-competent versus compromised styles was a Kunitz family protease inhibitor (Table 1). In *Nicotiana*, the Kunitz family protease inhibitor, NaStep, is highly expressed in the pistils of SI species and is thought to stabilize HT-proteins, although the mechanistic basis of this interaction remains unknown [67]. The NaStep protein is taken up by both compatible and incompatible pollen tubes, and the transgenic suppression of *NaStep* in SI *Nicotiana* species compromises rejection of both self-and some types of interspecific pollen tubes [67]. The reduced expression of this gene in UI-compromised LA0407 may reflect this population’s lack of ability to reject some types of interspecific pollen tubes [1, 6].

Although we were most interested in genes showing the highest upregulation in UI-competent styles, we also analyzed genes that are highly downregulated. We found that three of the top 25 most highly downregulated genes in UI-competent styles were involved in oxidation-reduction reactions, and one was putatively involved in defense response (Additional File 1, Table S5). Notably, five of the top 25 genes downregulated in UI-competent styles are putatively involved in cell wall modification (Additional File 1, Table S5). These include two Pectin Lyases (Sopen12g009500 and Sopen01g038750), a Glucosyltransferase (Sopen00g008620) and two Glycosyl Hydrolases (Sopen04g034210 and Sopen07g001240), all of which were upregulated over 13-fold. Eight additional genes involved in cell wall modification were downregulated in UI-competent styles, although to lesser extents (Additional File 1, Table S5).

In addition to our analysis of genes that are highly up-or downregulated in UI-competent versus compromised styles, we investigated the expression of 33 *a priori* candidates (Additional File 1, Table S6) based on reports in the literature of stylar genes that may be involved in UI [26, 28, 31, 54]. These candidates included genes that were identified in a recent study comparing stylar transcriptomes between UI-competent *S. pennellii* 0716 and the cultivated tomato *S. lycopersicum* M82 which shows no UI response [54]. Only three of the 33 *a priori* candidates showed over a 2-fold increase in styles of UI-competent LA1777 versus UI-compromised LA0407 (Additional File 1, Table S6). These included a Pectin-Methylesterase Inhibitor (Sopen04g027820, upregulated 2.4-fold) and two Glucosyltransferases (Sopen08g002330 and Sopen08g002350, both upregulated over 8-fold); however all were expressed at low levels, < 1 cpm. HT-protein, a small asparagine-rich protein required for S-RNase-based UI [33] was highly expressed in styles from both populations and was downregulated slightly in LA1777 (HT-A, 1.3-fold). Interestingly HT-B transcript was expressed in both populations, although neither population accumulates functional protein due to an early stop codon in the transcript [31]. Each individual of LA1777 harbored two unique S-RNase alleles as expected (data not shown), whereas LA0407 did not express S-RNase transcript, as has been documented in previous studies [31].

### Differential gene expression related to UI-competence in pollen

As coordinated interactions between pollen and pistil are required for successful fertilization, we also compared pollen transcriptomes between populations. Pollen tubes of LA1777 reach ovaries of all SI and SC species within the tomato clade [1]. However, LA0407 pollen tubes are rejected by all SI species, including by SI *S. habrochaites* LA1777 [1]. The inability of LA0407 pollen to traverse the styles of LA1777 and other SI *Solanum* species suggests that it has lost a pollen factor(s) required for S-RNase resistance. For our analysis, we therefore considered LA1777 pollen to be UI-competent, and LA0407 pollen to be UI-compromised.

We identified 90 genes that were upregulated in UI-competent pollen (Additional File 1, Table S7) and 99 that were downregulated (Additional File 1, Table S8). Ten of the top 25 genes upregulated in UI-competent versus UI-compromised pollen were annotated as hypothetical proteins (Table 2; Additional File 1, Table S7). Upon further analysis, the hypothetical gene with the highest upregulation in UI-competent pollen (Sopen12g014190) likely encodes an arabinogalactan protein (AGP) (Additional File 2, Fig. S2). In *Arabidopsis*, pollen AGPs are required for proper pollen tube development and growth [68–70], and pistil AGPs are known to stimulate pollen tube growth [69–72]. We also identified a Rab-GTPase (Rab4A, Sopen01g033860) that was upregulated nearly 200-fold in UI-competent pollen. Pollen specific proteins from this family have been found to promote pollen tube tip growth and play a role in the ability of the pollen tube to sense directional cues [73, 74].

**Table 2.** Top-25 upregulated genes in pollen of LA1777 vs. LA0407.

| Gene | p value | Mean expression | Fold change | Annotation |
| --- | --- | --- | --- | --- |
| Sopen12g014190.1 | 9.3E-05 | 26.2 | 2922.0 | [HP] Putative GPI-anchored AGP |
| Sopen01g010170.1 | 4.1E-04 | 3.7 | 1138.1 | Hypothetical protein |
| Sopen12g018710.1 | 1.1E-05 | 2.7 | 598.0 | KH Domain RNA-Binding Protein |
| Sopen06g022970.1 | 7.5E-05 | 2.7 | 591.3 | Isoflavone-7-O-Methyltransferase 9; flavonol biosynthesis |
| Sopen05g003280.1 | 3.0E-04 | 5.2 | 397.0 | F-box Protein CPR1/30; negative regulator of defense response |
| Sopen09g012310.1 | 7.7E-04 | 1.9 | 318.1 | Ribonuclease H-like superfamily protein; possible non-LTR retrotransposon family (LINE) |
| Sopen10g022570.1 | 2.7E-05 | 1.9 | 287.3 | Hypothetical protein |
| Sopen08g010210.1 | 7.1E-04 | 1.8 | 277.0 | [HP] Possible retroelement |
| Sopen01g033860.1 | 2.2E-04 | 1.5 | 198.4 | RabA4 subfamily of Rab GTPases; promotes tip growth of pollen tubes |
| Sopen01g032940.1 | 3.6E-04 | 1.5 | 189.8 | Aspartyl Protease family protein, CDR1-like |
| Sopen12g018720.1 | 2.0E-04 | 1.4 | 168.0 | Hypothetical protein |
| Sopen12g022970.1 | 6.9E-04 | 1.3 | 154.6 | F-box/RNI superfamily; plant-specific FBD domain |
| Sopen08g010180.1 | 6.2E-04 | 1.2 | 130.0 | Hypothetical protein |
| Sopen03g005740.1 | 1.5E-04 | 1.1 | 105.8 | Acyl Activating Enzyme, Benzoate-CoA Ligase |
| Sopen03g020850.1 | 5.5E-04 | 1.1 | 98.6 | Possible transcription factor; VND-interacting 1, NAC domain |
| Sopen12g027860.1 | 5.6E-04 | 1.0 | 86.2 | [HP] possible non-coding RNA |
| Sopen05g018880.1 | 1.6E-04 | 1.5 | 84.7 | Hypothetical protein |
| Sopen10g022630.1 | 7.8E-04 | 0.9 | 74.6 | [HP] Non-coding RNA |
| Sopen10g023190.1 | 2.1E-04 | 0.9 | 71.5 | [HP] Possible LTR retrotransposon |
| Sopen06g021510.1 | 5.4E-05 | 0.9 | 71.4 | Zeatin O-Glucosyltransferase-Like |
| Sopen12g006170.1 | 4.6E-06 | 9.5 | 70.0 | Basic Leucine Zipper 34-Like, transcription factor |
| Sopen03g007850.1 | 1.7E-04 | 0.8 | 57.8 | Tetratricopeptide Repeat (TPR)-Like superfamily protein; pre-mRNA splicing factor SYF1 |
| Sopen10g028190.1 | 3.2E-04 | 1.1 | 52.6 | Wall-Associated Receptor Kinase (WAK2-like), Ca<sup>2+</sup> binding, cell wall expansion |
| Sopen02g013470.1 | 6.6E-04 | 1.1 | 50.4 | Cysteine-Rich RLK |
| Sopen10g022640.1 | 5.4E-04 | 0.8 | 48.8 | [HP] Non-LTR retrotransposon family (LINE) |

Gene annotation from Spenn\_v2.0 was enhanced, when possible, for genes noted as hypothetical proteins (marked as [HP]).

Our analysis of pollen also identified differentially expressed genes that may be involved in protein degradation pathways (Table 2; Additional File 1, Table S7 and Table S8). These genes are of interest, as components of a pollen SCF (SKP-Cullin-F-box) E3 ubiquitin ligase complex have been implicated in both SI and UI, and are potentially involved in detoxifying pistil side factors such as S-RNases through the proteasomal degradation pathway [11, 15, 21, 26, 27, 29, 75]. We found two F-box/Skp-2 –like genes (Sopen05g003280 and Sopen12g022970) that were upregulated over 150-fold in UI-competent pollen (Table 2). Another gene involved in protein degradation, an aspartyl protease (Sopen01g032940) was highly upregulated 189-fold in UI-competent pollen (Table 2). No previously identified SLFs [28] were significantly differentially regulated, nor was the pollen UI factor Cullin1 [26] (Additional File 1, Table S9).

We identified two protein kinases that were upregulated over 50-fold in UI-competent pollen, one of which encodes a calcium binding serine/threonine wall-associated kinase (Sopen10g028190), and the other a cysteine-rich receptor like kinase (RLK) (Sopen02g013470) (Table 2). Because RLKs have a proven role in pollen tube growth [76], these genes are of potential interest. Further, pollen-expressed RLKs are involved in reception of peptide and hormone signals from the pistil [77–79].

Genes involved in transcriptional regulation are likely to be important components of pollen tube growth. We identified two types of transcription factors known to play key roles in stress response [80, 81] that were highly upregulated in UI-competent pollen: a NAC-domain containing protein (Sopen03g020850) and a bZIP family protein (Sopen12g006170) that has previously been localized to pollen (Table 2; Additional File 1, Table S7).

An analysis of genes that were highly downregulated in UI-competent pollen identified an F-box Protein (Sopen11g004020) and a Cysteine-rich RLK (Sopen05g014070) that were downregulated over 45-and 100-fold respectively (Supplemental Table S8). Genes showing over 100-fold decreases in UI-competent pollen also included two Histone Deacetylases (HDACs, Sopen11g004040 and Sopen11g004050; Additional File 1, Table S8). The proper function of HDACs has been linked to successful pollen tube germination and tip growth in *Picea willsoni* [82].

### UI competence in additional S. habrochaites populations

Our initial analyses comparing transcriptomes of LA1777 and LA0407 reproductive tissues led to some intriguing candidates that might be integral to UI. In an attempt to further narrow down candidate gene and to expand our analysis to a broader range of *S. habrochaites* populations, we performed a second RNA-seq experiment that included UP styles and pollen from *S. habrochaites* populations with selected phenotypes (Table 3; Additional File 1, Table S10).

**Table 3.** *Solanum habrochaites* populations used in this study.

| Country | Province or Department | Population/<br>Accession<sup>a</sup> | Mating<br>system<sup>b</sup> | Interpopulation UI<sup>c</sup> |  | Interspecific UI<sup>c</sup> |  |
| --- | --- | --- | --- | --- | --- | --- | --- |
|  |  |  |  | Pollen<br>accepted<br>by LA1777 | Accepts<br>pollen of<br>LA0407 | Accepts<br>pollen of<br>*S. neorickii* | Accepts<br>pollen of *S.*<br>*lycopersicum* |
| Ecuador | Guayas | **LA0407** | SC | I | SC | C | I |
| Ecuador | Chimborazo | **LA1223** | SC | I* | C | C | C |
| Ecuador | Chimborazo | **LA1264** | SC | C | C | I | I |
| Ecuador | Loja | **LA2119** | SC | C | C | I | I |
| Ecuador | Loja | **LA2098** | SI/SC | C | I | I | I |
| Peru | Ancash | **LA1777** | SI | SI | I | I | I |

<sup>a</sup> all populations and passport information was acquired from the Tomato Genetics Resource Center (TGRC, tgrc.ucdavis.edu, [103]) at the University of California, Davis.
<sup>b</sup> mating system was verified experimentally, and is consistent with data from the TGRC, Rick *et al.* 1979 and Broz *et al.* 2016 [6, 44].
<sup>c</sup> interpopulation and interspecific UI as reported in Broz *et al.*, 2016 [6].
\*pollen of the LA1223 individual used in this experiment was not accepted by LA1777, Broz *et al.*, 2016 [6] reports variation in individual LA1223 phenotypes for this cross SC, self-compatible; SI, self-incompatible; C, compatible; I, incompatible

First we confirmed that the UI phenotypes of the individuals tested reflected previous results [6] (phenotypes summarized in Table 3). A PCA of all samples shows that much of the sample variation (1^st^ PC) can be explained by tissue type, whereas a smaller percentage of variation (2^nd^ PC) is explained by source population and potentially mating system (Fig. 3). Interestingly, the additional populations selected for transcriptome analysis, which lie between LA1777 and LA0407 geographically, cluster between these two populations in the PCA.

**Figure 3.**
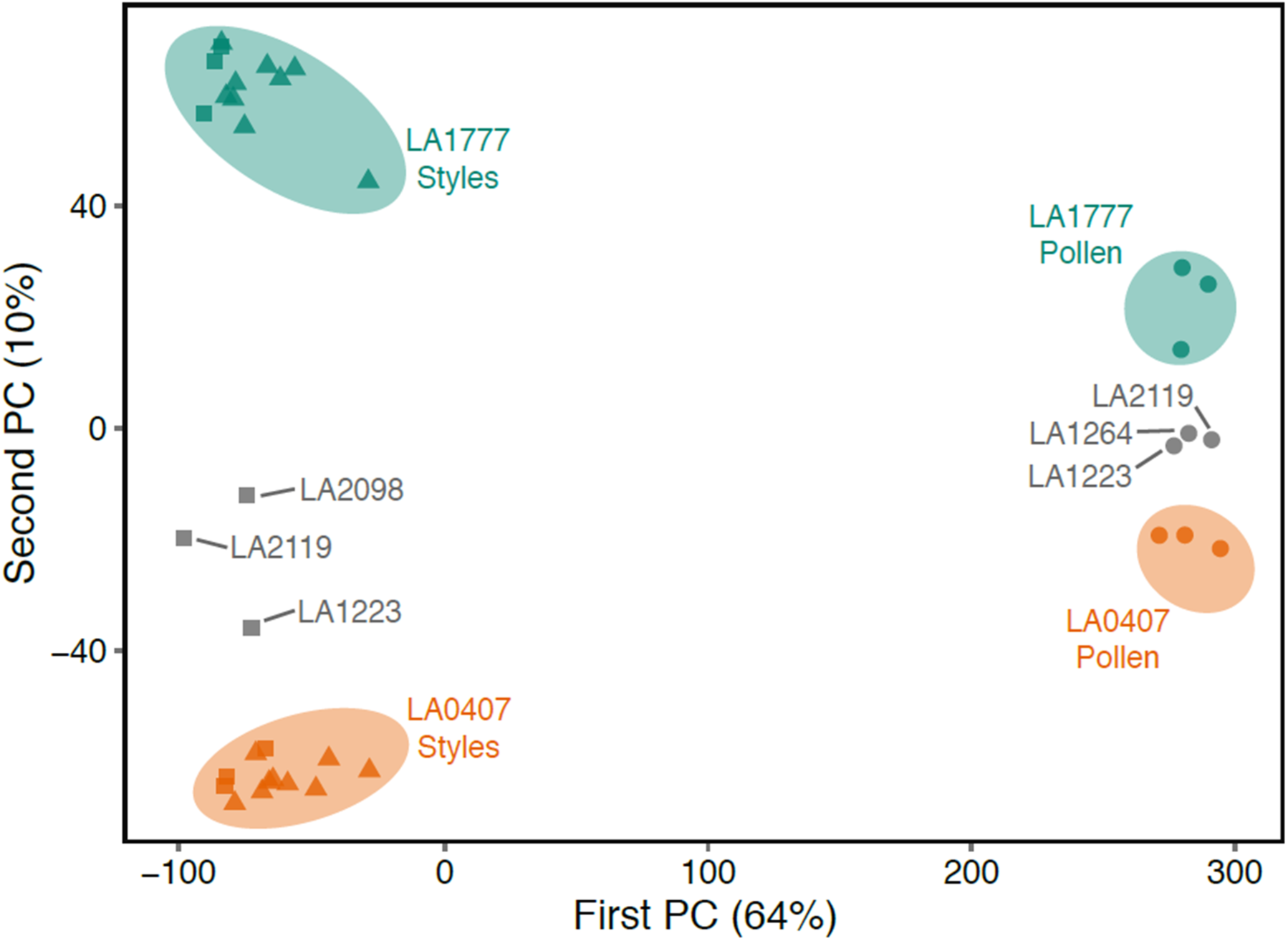
Genome-wide variation in gene expression across five populations of *Solanum habrochaites*, summarized by their first two principal components. The largest source of variation results from differences in expression between pollen (circles) and styles (unpollinated, squares; pollinated, triangles), on the first PC (horizontal axis). Variation across populations, on the other hand, accumulates on the second PC (vertical axis).

For each gene, we trained a linear discriminant function for UI-competence on the expression values for LA1777 and LA0407, and then classified gene expression in other populations as UI-competent or UI-compromised. We took into account the UI patterns of these populations with LA0407, as well as previously reported results on interspecific UI ([6], summarized in Table 3). Because styles of LA2119 are unique in that they accept interpopulation LA0407 pollen while rejecting interspecific pollen, we interpret the information in Table 4 to reflect the variation in UI phenotype of LA2119 styles (i.e., genes that are up in LA2119 are top candidates in interspecific UI, whereas genes that are down in LA2119 are top candidates in interpopulation UI).

**Table 4.** Upregulated genes in styles of LA1777 vs. LA0407 that show concordant expression patterns.

| **Interspecific UI (upregulated in LA2119)** |  |  |  |  |
| --- | --- | --- | --- | --- |
| **Gene** | **p value** | **Mean expression** | **Fold change** | **Annotation** |
| Sopen01g011020.1 | 4.0E-21 | 75.9 | 108.4 | Hypothetical protein |
| Sopen12g021490.1 | 7.0E-18 | 10.2 | 17.4 | NAD(P)-linked Oxidoreductase |
| Sopen07g030750.1 | 1.0E-13 | 4.7 | 6.1 | Cytochrome P450 (CYP72A15) |
| Sopen06g026260.1 | 5.5E-17 | 39.4 | 4.7 | Oxidoreductase, involved in lipid metabolic process |
| Sopen05g028380.1 | 2.0E-08 | 7.9 | 3.6 | Heavy metal transport/detoxification superfamily protein |
| Sopen10g033870.1 | 1.4E-10 | 12.7 | 3.3 | RING/U-box superfamily protein |
| Sopen12g006800.1 | 3.2E-13 | 73.7 | 3.1 | Photosystem II, Light-harvesting Chlorophyll B-binding protein |
| Sopen06g019380.1 | 1.6E-14 | 108.4 | 3.1 | Photosystem I Reaction Center Subunit II |
| Sopen03g005470.1 | 1.2E-13 | 6.9 | 3.0 | Peroxisomal Membrane 22 kDa (Mpv17/PMP22) family protein |
| **Interpopulation UI (downregulated in LA2119)** |  |  |  |  |
| **Gene** | **p value** | **Mean expression** | **Fold change** | **Annotation** |
| Sopen10g001450.1 | 8.8E-18 | 3.2 | 13.4 | Unknown chloroplast stroma protein |
| Sopen01g031170.1 | 5.3E-06 | 9.9 | 11.0 | Transcription factor GRAS, DELLA-like protein |
| Sopen05g006860.1 | 7.4E-14 | 74.6 | 6.5 | Alpha-1,4-Glucan-Protein Synthase, involved in cell wall biogenesis |
| Sopen11g005210.1 | 6.8E-19 | 33.1 | 5.3 | Alpha/beta-Hydrolases superfamily protein, Serine Hydrolase |
| Sopen11g028080.1 | 7.3E-16 | 5.2 | 4.2 | DHBP, RibB-like; 3,4-Dihydroxyl-2-Butanone 4-Phosphate Synthase |
| Sopen08g025270.1 | 7.9E-12 | 54.7 | 4.1 | ABC-transporter-like, P-Glycoprotein 2 (PGP2) |
| Sopen03g006340.1 | 5.2E-13 | 19.1 | 3.6 | Unknown function (DUF827) |
| Sopen07g027150.1 | 3.4E-10 | 131.8 | 3.5 | Phosphoenolpyruvate Carboxylase (PEPC) |
| Sopen12g001030.1 | 1.1E-08 | 9.2 | 3.4 | RING/U-box superfamily protein |
| Sopen04g034310.1 | 1.4E-09 | 2.8 | 3.4 | HXXXD-type Acyl-transferase family protein |
| Sopen11g001350.1 | 2.6E-11 | 11.0 | 3.1 | Plastid-lipid Associated Protein PAP / fibrillin family protein |
| Sopen08g021970.1 | 2.8E-13 | 17.8 | 3.1 | Hemoglobin Protein 3 (GLB3) |
| Sopen12g006260.1 | 5.1E-13 | 3.9 | 3.1 | F-box/LRR Protein, MAX2 |

A linear discriminant function was trained on the expression values of LA1777 and LA0407, and then used to classify other accessions as UI or non-UI.

The top candidates for interspecific UI-competence in styles identified in the linear discriminant analysis (LDA) included the hypothetical protein Sopen01g011020, that is highly expressed and upregulated >100-fold (Table 4). Six genes potentially involved in oxidation-reduction reactions were identified including the NAD(P)-linked Oxidoreductase (Sopen12g021490), a Cytochrome P450 (Sopen07g030750), a Lipid Oxidoreductase (Sopen06g026260), a Heavy Metal Transport Protein (Sopen05g028380), a Peroxisomal Membrane Protein (Sopen03g005470), and the Photosystem I Reaction Center (a Ferredoxin Oxidoreductase, Sopen12g006800).

Top candidates for interpopulation UI-competence in styles identified in the LDA included two potentially involved in ROS signaling (Table 4). A DELLA-like transcription factor (Sopen01g0311700) which is responsive to gibberellins [83] and participates in ROS signaling by regulating ROS accumulation [84], was upregulated 11-fold. A Hemoglobin Protein (Sopen08g021970) containing redox active transition metals was upregulated 3-fold. Other interpopulation UI candidates included the cell wall synthesis protein Alpha-1,4-Glucan-Protein Synthase (Sopen05g006860), an Alpha/beta Hydrolase family protein (Sopen11g005210), a P-Glycoprotein ABC Transporter-like protein (Sopen08g025270) and an F-box-LRR Protein (Sopen12g006260).

The LDA of pollen genes was more straightforward in that LA2119 and LA1264 were expected to be UI-competent (i.e., similar to LA1777) whereas LA1223 is UI-compromised and may be missing pollen factor(s) required to traverse SI styles. As shown in Table 5, using the LDA we identified 22 genes that were upregulated in UI-competent pollen. Two of these encode F-box proteins that are both putatively involved in the ubiquitin ligase complex (Sopen03g040880 and Sopen01g027170) and were both upregulated over 25-fold. In addition, a RAPTOR/KOG kinase homolog that is associated with the CUL4 RING ubiquitin ligase complex (Sopen10g027930) was upregulated 12-fold in UI-competent pollen, and an annexin-like calcium binding protein (Sopen05g030530) putatively involved in calcium signaling and polysaccharide transport was upregulated over 4-fold. Two transcription factors were also identified in the analysis: Sopen12g027920 encodes a suppressor of FRI1 that may act to recruit histone H3 methyltransferases and Sopen11g020250 encodes a zinc finger transcription factor.

**Table 5.** Upregulated genes in pollen of LA1777 vs. LA0407 that show concordant expression patterns.

| Gene | p value | Mean expression | Fold change | Annotation |
| --- | --- | --- | --- | --- |
| Sopen05g018880.1 | 1.58E-04 | 1.5 | 84.7 | Hypothetical protein |
| Sopen03g040880.1 | 8.68E-06 | 0.7 | 36.0 | RNI-like, F-box, ubiquitin-protein ligase activity |
| Sopen01g027170.1 | 7.75E-05 | 0.6 | 28.5 | RNI-like, F-box, Skp2-like |
| Sopen02g038110.1 | 7.81E-06 | 0.6 | 24.9 | Cytochrome P450 724B1 |
| Sopen02g036410.1 | 1.18E-05 | 64.8 | 16.7 | Putative Hexokinase |
| Sopen12g026740.1 | 3.26E-04 | 114.7 | 11.9 | Hypothetical protein |
| Sopen10g027930.1 | 4.98E-04 | 5.0 | 11.9 | RAPTOR/KOG homolog located in CUL4 RING ubiquitin ligase complex |
| Sopen01g008220.1 | 1.05E-04 | 1.9 | 9.3 | Hypothetical protein |
| Sopen05g029540.1 | 2.80E-04 | 0.4 | 8.8 | RNA-directed DNA polymerase |
| Sopen01g028020.1 | 5.26E-05 | 3.4 | 7.3 | Hypothetical protein |
| Sopen12g027920.1 | 4.39E-05 | 0.3 | 7.0 | Zinc Finger Transcription Factor SUF4, involved in histone methylation |
| Sopen12g004180.1 | 3.29E-04 | 112.9 | 4.6 | Oligouridylate Binding Protein 1B |
| Sopen01g005900.1 | 1.36E-04 | 111.6 | 4.5 | Hypothetical protein |
| Sopen05g030530.1 | 3.09E-04 | 20.8 | 4.5 | Annexin, Ca<sup>2+</sup>-binding protein |
| Sopen03g024730.1 | 4.76E-05 | 5.5 | 3.9 | DNA Helicase |
| Sopen03g027800.1 | 2.34E-04 | 10.9 | 3.8 | Exostosin |
| Sopen02g038520.1 | 6.46E-04 | 247.3 | 3.7 | Lysine-Histidine Transporter (LHT1) |
| Sopen11g020250.1 | 4.01E-04 | 5397.5 | 3.7 | GATA Type Zinc Finger Transcription Factor |
| Sopen06g022340.1 | 4.85E-04 | 6.5 | 3.5 | Disease resistance/zinc finger/chromosome condensation-like region domain containing protein |
| Sopen07g015260.1 | 3.61E-04 | 108.5 | 3.2 | Vacuolar Proton ATPase Subunit VHA-a isoform 2 |
| Sopen11g021110.1 | 1.90E-04 | 129.8 | 3.1 | Protein of unknown function, DUF593 |
| Sopen06g009470.1 | 2.13E-04 | 0.8 | 3.1 | Hypothetical protein |

A linear discriminant function was trained on the expression values of LA1777 and LA0407, and then used to classify other accessions as UI or non-UI.

## Discussion

Although UI is widespread in plant families, the underlying molecular basis of this unidirectional reproductive barrier is not well understood. Here, using a transcriptomic approach, we identified genes in both pollen and stylar tissues that represent strong candidates for involvement in UI. Overall, our analyses identified a large number of differentially expressed genes that are involved in oxidation-reduction reactions and ROS signaling. Oxidative stress responses are involved in a variety of plant physiological processes, including reproduction [57, 59, 85–87]. The production of ROS in plants generally results in one of two outcomes: adaptation to stress or programmed cell death [85]. The balanced interplay between pollen and pistil ROS production can display either of these results: signaling and detoxification are required for successful fertilization (adaptation to stress) and can also function in the incompatible (SI) response (cell death of incompatible pollen tubes). For example, in the Papaveraceae family ROS are recognized as key regulators of programmed cell death in incompatible (self) pollen tubes [63, 88] and recent studies of Rosaceae family member *Pyrus pyrifolia* have demonstrated a link between ROS accumulation, Ca^2+^ signaling, calmodulin levels and actin filament depolymerization during self-pollen tube rejection [62, 89]. One of the first indications that ROS might be involved in pollen tube growth and cross-compatibilities in the Solanaceae came from histochemical staining in styles *Petunia hybrid,* which demonstrated that peroxidase activity is found in unpollinated styles and decreases following compatible pollinations, but remains high during incompatible self-pollinations [90]. This pattern is also observed in an analysis of cytochrome P450 (CYP51G1-Sc) in the wild potato *S. chacoense,* where in compatible pollinations, mRNA levels of CYP51G1-Sc declines, but in incompatible (self) pollinations, levels of this cytochrome remain stable.

In our pairwise comparisons investigating changes between UP and pollinated styles, we found increases in ROS pathway members in incompatible interpopulation pollinations (Additional File 1, Table S3) but not incompatible self-pollinations (Additional File 1, Table S1) or compatible intrapopulation pollinations (Additional File 1, Table S2). For instance, an H_2_O_2_ transporter was upregulated over 16-fold only in incompatible interpopulation pollinations. These transporters generally pump reactive H_2_O_2_ into the apoplast, an acidic environment with low numbers of ROS scavengers, resulting in oxidative stress [85]. A K^+^ channel and a Ca^2+^-binding annexin protein, both of which were annotated as being pollen-expressed were also upregulated in interpopulation interactions, as was a Rab-GTPase. All three of these proteins can play important roles in ROS generation and signaling. In pollen tubes, annexins may provide an important link between Ca^2+^, the membrane and the cytoskeleton [91]. Further, many annexins form Ca^2+^ channels which are vital not only for the oscillating Ca^2+^ influx associated with pollen tube tip growth, but also for facilitating ROS signaling, cell elongation and cell wall remodeling [85, 92]. Interestingly, SKOR K^+^ channels like the one identified in our analysis can also act as ROS-activated Ca^2+^ channels [92]. Small GTPases, including Rabs, increase NADP(H)-oxidase activity in a Ca^2+^-dependent manner [58, 93], and their proper function is required for pollen tube growth [58, 60, 73, 74, 94]. For example, the overexpression of both active and mutant forms of the RAB11 protein leads to the inhibition of pollen tube growth in *Nicotiana* [74] suggesting that the correct balance of multiple Rabs is required for effective pollen tube growth.

In UI-competent styles, ten of the top 25 most highly upregulated genes are putatively involved in ROS generation and/or signaling, including an NADP(H)-oxidase and multiple cytochrome P450s (Table 1). We also identified two highly upregulated gene candidates in the flavonoid pathway, which could produce flavonoid compounds to act as pro-or anti-oxidants (Table 1). In our subsequent analysis using transcriptome data from additional *S. habrochaites* populations, we found that six of the nine candidate stylar genes that may be involved in interspecific UI were ROS pathway genes (Table 4). Surprisingly, only two ROS-linked genes were upregulated in interpopulation UI-competent styles, one of which encodes a DELLA-like transcription factor that inhibits ROS accumulation and restrains cell expansion [83, 84]. In sum, these results suggest a dynamic interplay between pollen and pistil that must be held in a tight balance for pollen tubes to successfully grow through styles to reach the ovary.

The generation of ROS is linked to cell expansion, growth, cell wall cross linking and callose deposition [95]. Pollen tube walls consist of a number of polymers (callose, cellulose and pectin) that are highly cross-linked to each other; however the mechanisms by which cell wall modification is regulated remains unclear [96–98]. Studies using microarray analysis have found an upregulation of cell-wall modification genes in pollinated versus unpollinated styles [51]. However, few have specifically investigated specificity to the SI or UI response (but see Pease et al. [54]). Using electron microscopy, de Nettancourt et al [99] found differences in callose deposition between SI and UI crosses in *S. peruvianum* wherein SI crosses showed large levels of callose deposition at the pollen tube tips, but interspecific crosses did not [99]. Our pairwise comparisons in UP versus pollinated LA1777 styles did not reveal any genes involved in cell wall modification. However, we did identify a number of differentially expressed genes involved in cell wall modification in our larger analysis of stylar tissue (Additional File l, Table S4 and Table S5), most of which were highly downregulated in UI-competent styles.

One of the most intriguing gene candidates from our analysis of stylar tissue included a putative RALF peptide hormone (Sopen02g033850) that was upregulated over 22-fold in UI-competent styles. Peptide hormone signaling is involved in numerous processes during pollen-pistil interactions, from pollen hydration to fertilization [77]. A tomato RALF has been shown to reduce pollen tube growth during specific windows of development [66], and therefore a stylar-secreted RALF could play a role in the rejection of interspecific pollen tubes. Another interesting style-expressed candidate to pursue is the Kunitz-like Protease Inhibitor that was upregulated in UI-competent styles. The *Nicotiana* NaStep protein from this family is required for the rejection of some (but not all) interspecific pollen [67], and this protease inhibitor may play a similar role in *S. habrochaites* UI. Finally, the most highly-upregulated stylar gene identified in UI-competent styles encodes a putative prenylated heavy-metal binding protein (Additional File 2, Fig. S2), that may be worthy of further investigation. Other proteins of this type have been identified in a variety of tissues, and the few that have been characterized have been implicated in stress response [100].

In UI-competent pollen, we identified a highly upregulated putative AGP (Sopen12g014190) (Table 2) that contains a signal peptide, a hydrophobic C-terminal domain, eight dipeptides that are found in known AGPs and is composed of > 35% Pro/Ala/Ser/Thr(PAST) amino acids: a defining characteristic of AGPs (Additional File 2, Fig. S2; [70, 101]). In *Arabidopsis,* pollen AGPs are required for pollen tube growth, and may be involved in complex signaling cascades [68–70], and pollen AGPs have been localized to pollen tube tips in some species [102]. Another gene that warrants further study is a Rab-GTPase (Sopen01g033860) that was upregulated nearly 200-fold in UI-competent pollen. It will be interesting to see if increasing the expression of these genes in UI-compromised pollen is able to increase rates of pollen tube growth through SI styles.

### Conclusions

Our analyses revealed differentially expressed genes that may contribute to reproductive incompatibility between populations of *S. habrochaites*. This work represents an important first step in understanding how unilateral barriers might arise between populations, and how they are maintained during speciation. The variability in UI responses between *S. habrochaites* populations provides an exciting opportunity in which to further analyze these candidate genes and link them to specific UI phenotypes.

## Materials and Methods

### Solanum habrochaites *plant material and growth*

*Solanum habrochaites* (S. Knapp & D. M. Spooner) is a wild relative of tomato that demonstrates variability in mating system [41, 44], as well as interspecific [31, 35] and interpopulation [1, 6–8] cross-compatibilities. Seeds from the *S. habrochaites* accessions (referred to hereafter as populations) used in this study (Table 3) were acquired from the C.M. Rick Tomato Genetic Resource Center (TGRC) at University of California, Davis (www.tgrc.ucdavis.edu, [103]), sterilized according to recommendations from the TGRC, grown under greenhouse conditions as previously described [1] for approximately 3 weeks, and transplanted in covered outdoor agricultural field plots at Colorado State University. Plants used in the study were randomized within a single block, and remained large and healthy throughout the experiment. Specific populations were chosen based on reproductive characteristics as described in Broz et al., 2016 [6], see Table 3 for more information.

### Pollen tube growth phenotypes

Pollen tube growth through the style was assessed for self, intrapopulation and reciprocal interpopulation crosses of *S. habrochaites* individuals, although for LA0407, self and intrapopulation crosses consistently resulted in fruit-set, so pollen tube growth was not assessed for every individual. For all crosses, buds were emasculated 1 day prior to anthesis (day -1), hand-pollinated 24 hours later (day 0, budbreak), and harvested into fixative (1:3 acetoethanol) 48 h after pollination. For crosses between LA1777 (female) and LA0407 (male), styles were harvested at various time points after pollination (12, 24 and 48 h) to determine the time at which pollen tube rejection occurred. Pollinations were typically performed in the late afternoon and collected the following morning. However, pollen tube growth through styles was similar for pollinations performed in both the morning and the afternoon (data not shown).

Pollinations were covered with mesh bags to prevent unintended pollen deposition by pollinators. Pollen tube growth was assessed using fluorescence microscopy as previously described [31], and the length of styles and the point in the style at which no more than three pollen tubes passed were measured using ImageJ 1.47v (http://rsb.info.nih.gov/ij/; [104]).

### Tissue collection, RNA extraction and library construction

The primary RNA-seq experiment was performed to identify genes involved in interpopulation pollen tube rejection observed in crosses between LA1777 females and LA0407 males. Samples consisted of unpollinated styles, pollinated styles (self-pollination, intrapopulation pollination, or interpopulation pollination), and pollen (Fig. 1; Additional File 1, Table S10) – resulting in five treatment/tissue types for each individual. Three individuals from each population were used as biological replicates, resulting in a total of 30 libraries. In a follow-up experiment designed to narrow the list of genes involved in interpopulation interactions, RNA samples from between one and three individuals were pooled at an equimolar ratio before library creation (Additional File 1, Table S10).

For all style samples, flowers were emasculated and pollinated as described above for pollen tube growth experiments and harvested 16 hrs post-pollination (the approximate time at which pollen tube rejection occurred in interpopulation crosses). Unpollinated controls underwent the same treatment (emasculation at day −1), except they were left unpollinated. For each individual plant, approximately 30 styles were collected for each treatment and pooled before RNA extraction. To minimize variation due to environmental conditions, all treatments were conducted on the same days and harvested at approximately the same time of day. Styles (including stigmas) were harvested directly into RNALater (Qiagen) and stored at 4º C for one week, after which styles were blotted dry and immediately frozen at −80° C until processing. Approximately 100 mg of pollen was harvested from each individual plant and immediately frozen at −80° C.

Tissues were ground using the Tissue-lyser (Qiagen), RNA was extracted using the Qiagen RNeasy Plant mini-kit and brought to a final concentration of 70-200 ng/uL. A subset of RNA samples was checked for quality by agarose gel electrophoresis and visualization by ethidium bromide staining. Sample quality was further evaluated using the Agilent 2200 RNA TapeStation system before library creation. Stranded, paired-end libraries of total RNA were generated for each sample using Illumina Truseq Stranded mRNA sample preparation kits. Libraries were pooled and distributed evenly across two lanes of Illumina HiSeq™ 2000 (Illumina Inc., San Diego, CA, USA). RNA quality control, library preparation, and pooling were performed by the Indiana University Center for Genomics and Bioinformatics. The raw transcriptome data is available on the NCBI SRA database (BioProject SRP069274).

### RNA-seq read processing and mapping

Prior to mapping and assembly, reads were trimmed and filtered using the SHEAR program (http://www.github.com/jbpease/shear; [54]). Briefly, SHEAR first uses the Scythe algorithm (https://github.com/vsbuffalo/scythe; [105]) to remove adapters from the 3′ end, and then filters low quality reads (mean Q<10), reads with >7 ambiguous bases (N’s), reads < 50 bp, and repetitive reads with mutual information score > 0.5. The program then trims reads on both ends by removing low quality bases (Q<20), poly-A or poly-T runs of n ≥ 12, and ambiguous bases. The appearance of AGATC at the 3′ end was also removed as we presumed it was an adapter fragment. We removed both reads in a pair if either one of them failed the filters. On average, 2.9% of all reads failed to pass the filter (min 2.5%, max 3.4%). The full command and parameters used for SHEAR can be found in Additional File 2, Method S1.

We mapped RNA-seq reads to the *Solanum pennellii* reference genome using the STAR spliced aligner with default parameters [106]. The reference genome sequence and the genome annotation (Spenn v2.0) were downloaded from solgenomics.net [107]. On average across libraries, 81% of reads mapped uniquely to the reference genome. We counted reads mapped to the reference gene annotation and to unannotated putative S-locus F-box (SLF) genes [28] (a total of 48938 genic regions) using featureCounts v1.4.5-p1 [108]. A total of 663185500 read pairs were counted (72% of raw reads). For genes annotated as ‘hypothetical’ in Spenn v2.0 [107], further sequence alignments were carried out using NCBI BLAST searches (https://blast.ncbi.nlm.nih.gov; [109]).

### Differential gene expression analysis

For all tests of differential expression we used linear models implemented by the *limma* package [110, 111] and modules from the edgeR package [112] in R [113]. First, we normalized and transformed the raw reads with *voom,* a weighted transformation based on the expected relationship between expression mean and variance [111]. We computed *t*-statistics on the transformed expression values for each gene using an empirical Bayes adjustment of standard errors with the *eBayes* function [110].

We initially searched for differential expression among stylar tissues by carrying out separate pairwise comparisons. Specifically, we compared the following pairs of style treatments in LA1777: intrapopulation-pollinated (compatible) against unpollinated (UP) styles, self-pollinated (incompatible) against UP styles and interpopulation (incompatible) against UP styles.

We visualized the genome-wide patterns of expression through a PCA of the normalized mean read counts per gene (cpm reads in the library) with the *prcomp* function (implemented in R; [113]). The PCA showed that there were negligible differences at the genome-wide scale among all style treatments within a population (Fig. 3; Additional File 2, Fig. S1). Because of these high consistencies in their gene expression profiles, all style treatments (UP, self-, intra-and interpopulation pollinated) within a population were considered identical in our linear models that focus on the differences between UI-competent (LA1777) and UI-compromised (LA0407) styles.

We identified genes that are differentially expressed in UI-competent tissues using a linear model with a single fixed effect (collinear with the population of origin) for pollen (n=6) and styles (n = 24) separately. From these models, we took genes as differentially expressed if they showed large differences in expression (> 3-fold change), with statistical significance at a false discovery rate (FDR) of 5%. Further, we considered only tissue-specific genes: we required genes to have a significant (FDR < 5%) tissue effect in a general linear model *Y* ~ *P* + *T* + *e* (where *P* is a population effect, *T* is a tissue effect, and *e* is the error term). For stylar-side factors, we included genes as differentially expressed if they were upregulated in styles with respect to pollen, and vice versa for pollen-side factors. This last filter ensured that differences in styles were unlikely to be contributed by pollen in the styles of pollinated samples.

A number of *a priori* pollen and pistil UI candidate genes were selected based on information from previous publications [23, 26, 28, 31, 54]. Some of the SLF genes have not yet been annotated in the *S. pennellii* genome, so we used locus numbers and sequences from Li and Chetelat (2015) [28] to find these genes in our dataset. We identified expression levels of all *a priori* candidates, and carried out statistical tests similar to the ones described above to determine whether they were differentially expressed.

## Acknowledgements

The authors thank the Charles M. Rick Tomato Genetics Resource Center for seeds, J. Pease, M. Wu, and L. Moyle for assistance with the analysis, A. Ashford for plant care and O. Todd, T. Randall, and A. Martin for help compositing microscopic images. This work was supported by grant numbers DBI-0605200 and MCB-1127059 from the Plant Genome Research Program of the National Science Foundation.

## Authors’ contributions

AB performed RNA extraction, analyzed and interpreted data and wrote the manuscript with editing contributions from all authors. RG performed all bioinfomatics and statistical analyses of the data. AR designed and performed crossing experiments and collected tissue. YB performed crossing experiments and collected tissue. MH designed experiments, oversaw bioinformatics experiments and assisted in data analysis. PB conceived and designed experiments, collected tissue and interpreted data. All authors read and approved the manuscript.

---

**Figure S1.**
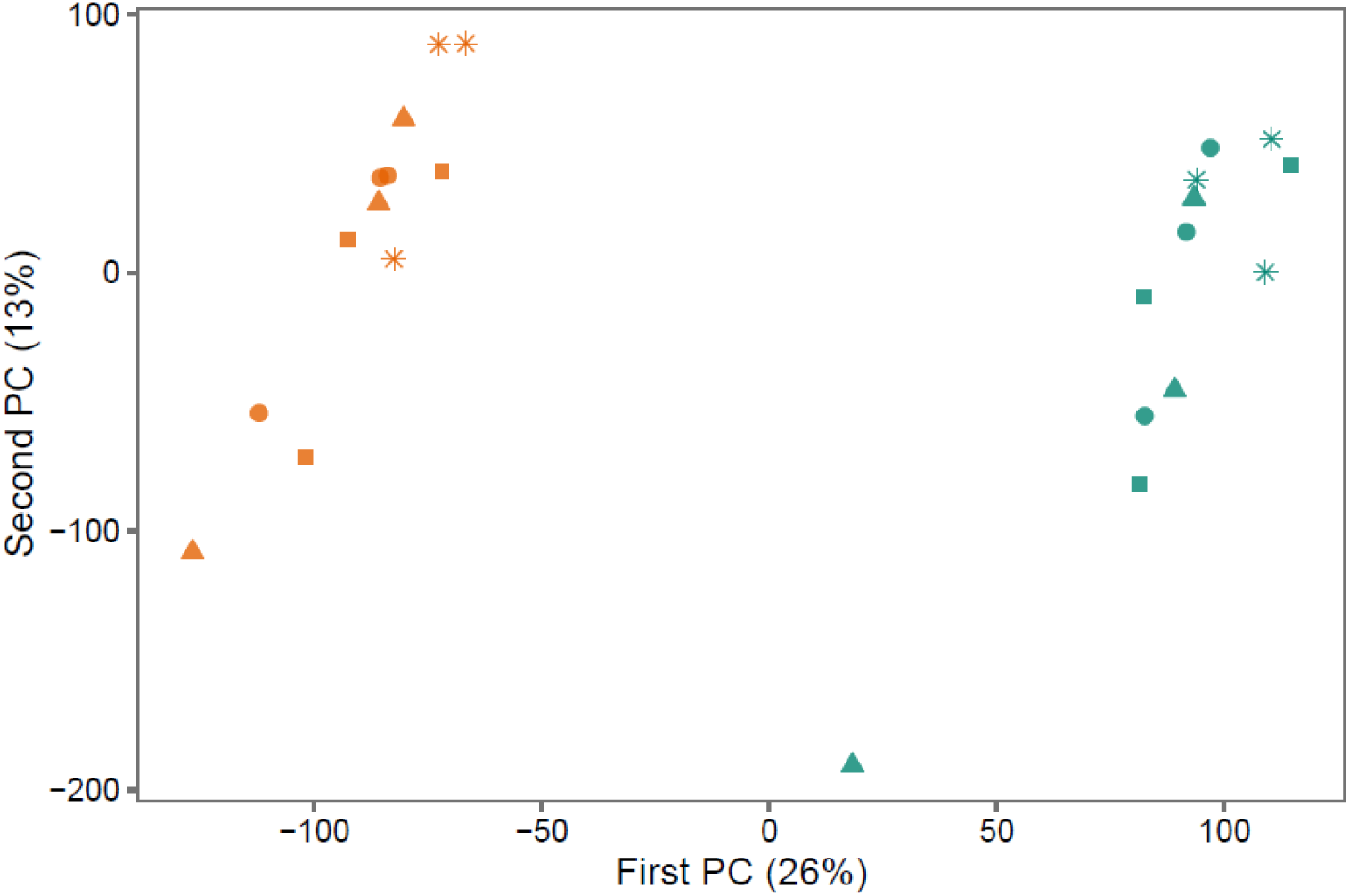
The genome-wide patterns of expression in styles from two populations of *Solanum habrochaites* (LA1777 in orange and LA0407 in green), summarized by a principal components analysis. Most variance among samples is due to differences between populations (horizontal axis), and we found no large differences among experimental treatments at this scale (unpollinated styles, stars; self-pollinated, circles; intrapopulation-pollinated, triangles; interpopulation pollinated, squares).

**Figure S2.**
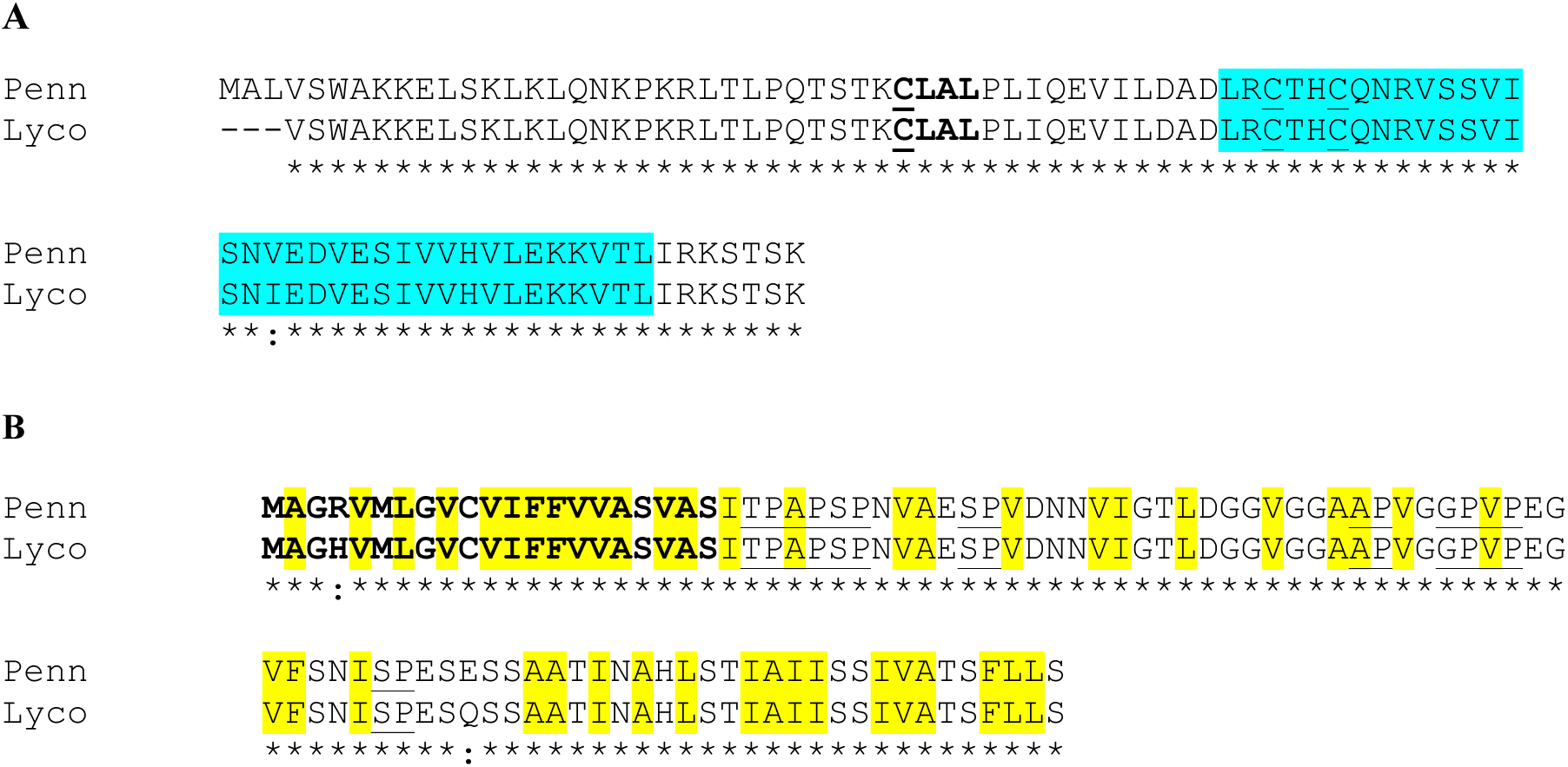
Sequence alignments of hypothetical proteins showing the highest fold-change in UI-competent vs. UI-compromised tissues. **A,** The deduced amino acid sequence of *Solanum pennellii* Sopen12g017530 (Penn), the gene showing the highest fold change in UI-competent versus UI-compromised styles, is aligned to a *Solanum lycopersicum* (Lyco) EST (GenBank # AW092656.1) identified in an elicitor screen of tomato leaf. A BLAST search of *S. lycopersicum* gene models (SOLv2.4, solgenomics.net) did not return any results. Putative heavy metal binding domain (P-Fam 00403) is shown in blue; putative prenylation site (CaaX, where ‘a’ represents an aliphatic amino acid) is shown in bold “CLAL”. Cysteine residues involved in both domains are underlined. Stars represent residues conserved between both sequences. **B,** The deduced amino acid sequence of *Solanum pennellii* Sopen12g014190 (Penn), the gene showing the highest fold change in UI-competent versus UI-compromised pollen, is aligned with *Solanum lycopersicum* Solyc12g033100 (Lyco). Putative signal peptide is shown in bold; distinguishing dipeptide motifs of arabinogalactan proteins (AGPs) are underlined. Hydrophobic residues are highlighted in yellow. Stars represent residues conserved between both sequences.

**Method S1.** Full command and parameters used for SHEAR in bioinformatics analysis of *S. habrochaites* transcriptomes.

~~~
#!/bin/bash
# These script snippets were used for the differential gene expression analysis presented in
# Broz et al. "Transcriptomic characterization of a pollen-pistil unilateral reproductive barrier"
# These will not run adequately without helper files (as indicated below)
# Software needed:
# scythe (github.com/vsbuffalo/scythe)
# shear (github.com/jbpease/shear)
# subread (subread.sourceforge.net/)
# STAR (github.com/alexdobin/STAR/)
# samtools (www.htslib.org/)
# Contact: Rafael F Guerrero, rafguerr@indiana.edu
# (1) Shear reads
#raw_R1_fullpaths.txt must contain all the names of the R1 fastq files to be analyzed
while read line
do
a=${line:0:${#1}-9}
python shear.py \
--fq1 $a"_R1.fastq" \
--fq2 $a"_R2.fastq" \
--out1 $a"_sheared_R1.fastq" \
--out2 $a"_sheared_R2.fastq" \
--execscythe /N/soft/rhel6/scythe/0.992beta/scythe \
--tempdir $(pwd)/tempfiles_shear \
--trimfixed 0:0 --trimqual 20:20 --trimqualpad 0:0 --filterlength 50 --
trimpattern3 AGATC --trimpolyat 12 --trimambig --filterlowinfo 0.5 --
filterunpaired --filterqual 10 --filterambig 8
done < raw_R1_fullpaths.txt
# (2) STAR mapping
#shear_R1_fullpaths.txt must contain all the names of the R1 fastq files
preprocessed by shear
# The variable PATH_TO_GENOME must be set to the absolute path to the (STAR
indexed) reference genome
while read line
do
prefix=${line:0:${#line}-17}
dirname=${prefix}_penn
if [ ! -d "$dirname"]; then
mkdir $dirname
cd $dirname
STAR \
--genomeDir $PATH_TO_GENOME \
--readFilesIn ${prefix}_sheared_R1.fastq /${prefix}_sheared_R2.fastq \
--runThreadN 8 \
--outReadsUnmapped Fastx \
--genomeLoad LoadAndKeep
fi
done < shear_R1_fullpaths.txt
# (3) Sorting, filtering and indexing alignment files
#dirnames.txt must have the absolute paths to the STAR output directories
(one per library)
while read line
do
cd $line
prefix="Aligned.out"
if [ ! -f ${prefix}".sorted.bam" ]; then
samtools view -uS ${prefix}".sam" > ${prefix}".bam"
samtools sort $prefix".bam" $prefix".sorted"
samtools index $prefix".sorted.bam"
rm $prefix".bam"
fi
done < dirnames.txt
# (4) Counting reads in gene models with featureCounts from subread-1.4.6
featureCounts -T 15 -B -p -t exon -g Parent -a spenn_v2.0_exons.gff -o
all_counts_to_PENN.txt $(cat all_penn_bams.txt)
~~~

